## Supplementary material for "*NCBP2* modulates neurodevelopmental defects of the 3q29 deletion in *Drosophila* and *X. laevis* models": S1 Table

| **Human gene** | **Fly homolog** | **Identity (%)** | **Similarity (%)** | **DIOPT score** | **DIOPT rank** | **Larval central nervous system expression** (FlyAtlas) | **Larval eye expression** (modENCODE) |
|---|---|---|---|---|---|---|---|
| *BDH1* | *CG8888* | 33 | 53 | 9 | High | Low | NA |
| *DLG1* | *dlg1* | 44 | 58 | 13 | High | Moderate | Moderate |
| *FBXO45* | *Fsn* | 71 | 84 | 13 | High | Moderate | Moderate |
| *MFI2* | *Tsf2* | 33 | 48 | 15 | High | Low | Moderate |
| *NCBP2* | *Cbp20* | 78 | 89 | 14 | High | Moderate | Moderate |
| *OSTalpha* | *CG6836* | 19 | 40 | 5 | High | Low | Low |
| *PAK2* | *Pak* | 42 | 50 | 10 | Moderate | NA | Moderate |
| *PCYT1A* | *Pcyt2* | 58 | 72 | 12 | High | Moderate | Moderate |
| *PIGX* | *PIG-X* | 24 | 39 | 7 | High | Low | Low |
| *PIGZ* | *PIG-Z* | 30 | 41 | 10 | High | NA | Low |
| *SENP5* | *Ulp1* | 21 | 35 | 2 | Low | Moderate | Low |
| *TCTEX1D2* | *CG5359* | 33 | 51 | 9 | Moderate | Moderate | Low |
| *UBXN7* | *CG8892* | 28 | 43 | 13 | High | Moderate | Moderate |
| *WDR53* | *CG5543* | 21 | 34 | NA | NA | Low | Moderate |
| *ZDHHC19* | *app* | 34 | 49 | 3 | Moderate | NA | Low |
| *CEP19* | None | | | | | | |
| *LRRC33* | None | | | | | | |
| *RNF68* | None | | | | | | |
| *SMCO1* | None | | | | | | |
| *TFRC* | None | | | | | | |
| *TM4SF19* | None | | | | | | |
