## Supplementary material for "*NCBP2* modulates neurodevelopmental defects of the 3q29 deletion in *Drosophila* and *X. laevis* models": S2 Table

| Gene name | Forward and reverse primers | RNAi knockdown (% expression) |
|---|---|---|
| *app*$^{KK108227}$ | For-5'- GCGATCAGACAACCAACGAG-3'<br>Rev-5'- CGCCTTTGGAGGAGAAGGAT-3' | 55.457 |
| *Cbp20*$^{KK109448}$ | For-5'- TTGTGAATGGCACTCGCTTG-3'<br>Rev-5'- GTCCCAGTCCACACGAATCA-3' | 43.900 |
| *CG5359*$^{KK107839}$ | For-5'- ACGTTATGGCCGAGAAACTCA-3'<br>Rev-5'-TGGCGACGTCCTTGTCATAG-3' | 20.945 |
| *CG5543*$^{KK109031}$ | For-5'- AAATCCACTTAGCGTGGGGC-3'<br>Rev-5'- AGGAAATTTTACCGCGTTGCAT-3' | 49.764 |
| *CG6836*$^{KK112485}$ | For-5'- CCCTTCATCGTCTGCTCCAT-3'<br>Rev-5'- GTGATTTGGAGGGACCAAGC-3' | 49.087 |
| *CG8888*$^{GD3777}$ | For-5'- TTCGCAAGAGCTTGGACCTC-3'<br>Rev-5'- TTTGTGTTAGCCGAGCGGAA-3' | 25.005 |
| *CG8892*$^{GD14061}$ | For-5'- TCCAGAGCAACGTCATGTCC-3'<br>Rev-5'- TGGACCGTCTGTTAAGTGCC-3' | 38.721 |
| *dlg1*$^{GD4689}$ | For-5'- ACACAAGACGATGCCAATGC-3'<br>Rev-5'- TCCACCCTGTAGATAATCTCGC-3' | 62.691 |
| *Fsn*$^{GD11383}$ | For-5'- CCCATTTGGTTGGTGTGGGA-3'<br>Rev-5'- TGGATTTACCCGTTCCTGTTGA-3' | 55.230 |
| *Pak*$^{KK101874}$ | For-5'- GTCGTCACCCGGAAACAGTA-3'<br>Rev-5'- GCCCAAAGACCAAAGGTCCA-3' | 40.212 |
| *Pcyt2*$^{KK110819}$ | For-5'- CGCTACGTGGATGAGATCGT-3'<br>Rev-5'- TCCTCATTTAGCGTCCACGG-3' | 80.642 |
| *PIG-X*$^{KK109717}$ | For-5'- TGACCTGCAGCGTTTGAAGA-3'<br>Rev-5'- TGACGAACTTAGGATAGATGGCA-3' | 29.775 |
| *PIG-Z*$^{KK107404}$ | For-5'-TCCAGAGCGTGGAGGTAATG-3'<br>Rev-5'- CGTATGCTCCAGCCGAAAGT-3' | 37.856 |
| *Ulp1*$^{GD7581}$ | For-5'-CCTGGCCAAGGGCTAAAAGT-3'<br>Rev-5'- GACATGCGTGTGTTCGCTAC-3' | 30.077 |
| *Rp49* control | For-5'-GCAAGCCCAAGGGTATCGA-3'<br>Rev-5'-ACCGATGTTGGGCATCAGA-3' | -- |
| *tiptop* | For-5'-CCTCCACAGCATCAGCAACA-3'<br>Rev-5'-CCACCAGGTCGTTACCGTTC-3' | -- |
