## Supplementary material for "*NCBP2* modulates neurodevelopmental defects of the 3q29 deletion in *Drosophila* and *X. laevis* models": S3 Table

| 3q29 genes | *Drosophila* phenotypes | | | | | *X. laevis* phenotypes | | | | | | | Mouse phenotypes | | | | | | |
|---|---|---|---|---|---|---|---|---|---|---|---|---|---|---|---|---|---|---|---|
| | Behavior | Development | Lethality | Neuroanatomy | Neurophysiology | Behavioral/ neurological | Cellular | Embryo | Growth/size/body | Mortality/aging | Nervous system | Other | Behavioral/ neurological | Cellular | Embryo | Growth/size/body | Mortality/aging | Nervous system | Other |
| *BDH1 (CG8888)* | X | X | | X | | | | | | | | | | | | | | | |
| *CEP19* | | | | | | | | | | | | | | | | | | | |
| *DLG1 (dlg1)* | X | | X | X | | | | | | | | | | | | | | | |
| *FBXO45 (Fsn)* | | | | | | | X | | | | X | X | | | | | | | |
| *LRRC33* | | | | | | | | | | | | | | | | | | | |
| *MFI2 (Tsf2)* | | X | X | | | | | | | | | | | | | | | | |
| *NCBP2 (Cbp20)* | X | X | X | X | | | X | | | | X | X | | | | | | | |
| *OSTalpha (CG6836)* | X | | X | | | | | | | | | | | | | | | | |
| *PAK2 (Pak)* | X | X | | X | | | | | | | X | X | | | | | | | |
| *PCYT1A (Pcyt2)* | | | | | | | | | | | | | | | | | | | |
| *PIGX (PIG-X)* | X | X | | X | | | | | | | | | | | | | | | |
| *PIGZ (PIG-Z)* | X | | | X | | | | | | | | | | | | | | | |
| *RNF168* | | | | | | | | | | | | | | | | | | | |
| *SENP5 (Ulp1)* | | | X | | | | | | | | | | | | | | | | |
| *SMCO1* | | | | | | | | | | | | | | | | | | | |
| *TCTEX1D2 (CG5359)* | | | | | | | | | | | | | | | | | | | |
| *TFRC* | | | | | | | | | | | | | | | | | | | |
| *TM4SF19* | | | | | | | | | | | | | | | | | | | |
| *UBXN7 (CG8892)* | | | | | | | | | | | | | | | | | | | |
| *WDR53 (CG5543)* | | | X | X | | | | | | | | | | | | | | | |
| *ZDHHC19 (app)* | X | | | X | | | | | | | | | | | | | | | |
