## Supplementary material for "*NCBP2* modulates neurodevelopmental defects of the 3q29 deletion in *Drosophila* and *X. laevis* models": S4 Table

| **RNAi line** | **Mild axon targeting phenotypes** | **Moderate axon targeting phenotypes** | **Severe axon targeting phenotypes** |
|---|---|---|---|
| *Cbp20*$^{KK109448}$ | 4/9 | 3/9 | 2/9 |
| *dlg1*$^{GD4689}$ | 0/7 | 2/7 | 5/7 |
| *Fsn*$^{GD11383}$ | 7/20 | 7/20 | 6/20 |
| *Pak*$^{KK101874}$ | 2/8 | 4/8 | 2/8 |
| *Cbp20*$^{KK109448}$/ *dlg1*$^{GD4689}$ | 2/17 | 8/17 | 7/17 |
| *Cbp20*$^{KK109448}$/ *Fsn*$^{GD11383}$ | 1/16 | 4/16 | 11/16 |
| *Cbp20*$^{KK109448}$/ *Overexp. Diap1* | 5/11 | 6/11 | 0/11 |
| *dlg1*$^{GD4689}$/ *Overexp. Diap1* | 1/17 | 8/17 | 8/17 |
