## Supplementary material for "*NCBP2* modulates neurodevelopmental defects of the 3q29 deletion in *Drosophila* and *X. laevis* models": S5 Table

| Fly RNAi line | Human homolog | CNV region | Avg. eye phenotypic score |
|---|---|---|---|
| *Ube3a*$^{KK104898}$ | *UBE3A* | Core gene | 59.733 |
| *Pten*$^{GD13500}$ | *PTEN* | Core gene | 58.275 |
| *Cadps*$^{GD9502\_1}$ | *CADPS2* | Core gene | 56.758 |
| *PIG-Z*$^{KK107404}$ | *PIGZ* | 3q29 | 56.243 |
| *arm*$^{KK102545}$ | *CTNNB1* | Core gene | 54.865 |
| *app*$^{KK108227}$ | *ZDHHC19* | 3q29 | 53.614 |
| *kis*$^{GD16331}$ | *CHD8* | Core gene | 51.182 |
| *Nrx-1*$^{GD2619}$ | *NRXN1* | Core gene | 48.753 |
| *Prosap*$^{GD10101}$ | *SHANK3* | Core gene | 48.748 |
| *Cbp20*$^{KK109448}$ | *NCBP2* | 3q29 | 46.268 |
| *dlg1*$^{GD4689}$ | *DLG1* | 3q29 | 43.219 |
| *CG5543*$^{KK109031}$ | *WDR53* | 3q29 | 40.349 |
| *CG8888*$^{GD3777}$ | *BDH1* | 3q29 | 39.126 |
| *rk*$^{GD14383\_1}$ | *LGR5* | Core gene | 38.021 |
| *MCPH1*$^{GD12537\_2}$ | *MCPH1* | Core gene | 36.835 |
| *Pak*$^{KK101874}$ | *PAK2* | 3q29 | 36.691 |
| *para*$^{GD3392\_1}$ | *SCN1A* | Core gene | 35.846 |
| *PIG-X*$^{KK109717}$ | *PIGX* | 3q29 | 34.392 |
| *Eph*$^{GD39}$ | *EPHA6* | Core gene | 31.468 |
| *CG8892*$^{GD14061}$ | *UBXN7* | 3q29 | 31.179 |
| *CG6836*$^{KK112485}$ | *OSTalpha* | 3q29 | 30.842 |
| *Ulp1*$^{GD7581}$ | *SENP5* | 3q29 | 30.383 |
| *Pcyt2*$^{KK110819}$ | *PCYT1A* | 3q29 | 28.423 |
| *Fsn*$^{GD11383}$ | *FBXO45* | 3q29 | 27.671 |
