## Supplementary material for "*NCBP2* modulates neurodevelopmental defects of the 3q29 deletion in *Drosophila* and *X. laevis* models": S6 Table

| RNAi line | Cone cell defect | Primary cell defect | Secondary cell defect | Bristle group defect | Rotation error | Hexagonal defect | Photoreceptor defect |
|---|---|---|---|---|---|---|---|
| *GMR-GAL4* | | | | | | | |
| *Cbp20*$^{KK109448}$ | ++ | + | ++ | ++ | ++ | ++ | +++ |
| *CG5543*$^{KK109031}$ | ++ | ++ | ++ | +++ | ++ | + | + |
| *CG6836*$^{KK112485}$ | + | | | + | + | | + |
| *CG8888*$^{GD3777}$ | ++ | | | + | ++ | | ++ |
| *CG8892*$^{GD14061}$ | | | | | | | + |
| *dlg1*$^{GD4689}$ | ++ | | + | +++ | + | ++ | +++ |
| *Fsn*$^{GD11383}$ | ++ | + | | ++ | ++ | + | |
| *Pak*$^{KK101874}$ | + | | | + | + | + | |
| *Pcyt2*$^{KK110819}$ | + | ++ | ++ | ++ | ++ | + | + |
| *PIG-X*$^{KK109717}$ | + | | + | ++ | ++ | | + |
| *PIG-Z*$^{KK107404}$ | + | | + | ++ | + | + | ++ |
| *Ulp1*$^{GD7581}$ | + | ++ | ++ | + | ++ | + | + |
