## Supplementary material for "*NCBP2* modulates neurodevelopmental defects of the 3q29 deletion in *Drosophila* and *X. laevis* models": S7 Table

| **Second-hit homolog** | *app*$^{KK108227}$ | *Cbp20*$^{KK109448}$ | *CG6836*$^{KK112485}$ | *CG8888*$^{GD3777}$ | *dlg1*$^{GD4689}$ | *Fsn*$^{GD11383}$ | *Pak*$^{KK101874}$ | *PIG-X*$^{KK109717}$ | *PIG-Z*$^{KK107404}$ |
|---|---|---|---|---|---|---|---|---|---|
| *app* | NA | Enhancer (1/1) | No interaction (0/1) | No interaction (0/1) | Enhancer (1/1) | NA | No interaction (0/1) | No interaction (0/1) | No interaction (0/1) |
| *Cbp20* | Enhancer (1/1) | Enhancer (3/3) | Enhancer (1/1) | Enhancer (3/3) | Enhancer (2/2) | Enhancer (2/3) | Enhancer (3/3) | No interaction (0/1) | Enhancer (3/3) |
| *CG6836* | No interaction (0/1) | Enhancer (1/1) | NA | Enhancer (1/1) | Enhancer (1/1) | No interaction (0/1) | No interaction (0/1) | No interaction (0/1) | No interaction (0/1) |
| *CG8888* | No interaction (0/1) | Enhancer (3/3) | Enhancer (1/1) | Not validated (1/2) | Not validated (1/3) | Enhancer (2/3) | No interaction (0/3) | No interaction (0/1) | Enhancer (3/3) |
| *dlg1* | Enhancer (1/1) | Enhancer (4/4) | No interaction (0/1) | Not validated (1/2) | Enhancer (1/1) | Not validated (1/2) | Not validated (1/2) | No interaction (0/1) | Enhancer (3/3) |
| *Fsn* | Enhancer (1/1) | Enhancer (3/3) | No interaction (0/1) | Not validated (1/3) | Not validated (1/3) | No interaction (0/2) | No interaction (0/2) | No interaction (0/1) | Enhancer (2/3) |
| *Pak* | No interaction (0/1) | Enhancer (3/3) | No interaction (0/1) | Not validated (1/3) | Not validated (1/3) | No interaction (0/1) | No interaction (0/1) | No interaction (0/1) | Enhancer (2/3) |
| *PIG-X* | No interaction (0/1) | Enhancer (1/1) | No interaction (0/1) | No interaction (0/1) | No interaction (0/1) | No interaction (0/1) | No interaction (0/1) | NA | No interaction (0/1) |
| *PIG-Z* | No interaction (0/1) | Enhancer (2/2) | No interaction (0/1) | Enhancer (2/2) | Not validated (1/2) | Not validated (1/2) | No interaction (0/2) | No interaction (0/1) | Enhancer (1/1) |
| *CG5543* | NA | Enhancer (2/2) | NA | Enhancer (2/2) | Not validated (1/2) | NA | Not validated (1/2) | NA | NA |
| *CG8892* | NA | Enhancer (1/1) | NA | No interaction (0/1) | Enhancer (1/1) | NA | No interaction (0/1) | NA | NA |
| *Pcyt2* | NA | Enhancer (1/1) | NA | Enhancer (1/1) | No interaction (0/1) | NA | Enhancer (1/1) | NA | Enhancer (1/1) |
| *Tsf2* | NA | Enhancer (1/1) | NA | Enhancer (1/1) | No interaction (0/1) | No interaction (0/1) | Enhancer (1/1) | NA | Enhancer (1/1) |
| *Ulp1* | NA | No interaction (0/2) | NA | Not validated (1/2) | Not validated (1/2) | NA | Not validated (1/2) | NA | NA |
| Lines tested **(161 total)** | 8 | 28 | 8 | 25 | 24 | 16 | 23 | 8 | 21 |
| All interactions **(54/94 total)** | 3/8 | 12/13 | 2/8 | 10/13 | 10/13 | 4/8 | 6/13 | 0/8 | 7/10 |
| Validated **(39/94 total)** | 3/8 | 12/13 | 2/8 | 6/13 | 4/13 | 2/8 | 3/13 | 0/8 | 7/10 |
| Reciprocal cross **(19/26 total)** | 2/2 | 7/8 | 2/2 | 3/3 | 1/3 | 1/2 | 1/1 | 0/0 | 2/5 |
