## Supplementary material for "*NCBP2* modulates neurodevelopmental defects of the 3q29 deletion in *Drosophila* and *X. laevis* models": S8 Table

| Pairwise cross | Cone cell defects | | | Primary cell defect | Secondary cell defect | Bristle cell defect | Rotation error | Hexagonal defect | Photoreceptor defect |
|---|---|---|---|---|---|---|---|---|---|
| | Number error | Arrangement error | Orientation error | | | | | | |
| *dlg1*$^{GD4689}$ | | | ++ | | + | +++ | + | ++ | +++ |
| *Cbp20*$^{KK109448}$ | | | ++ | + | ++ | ++ | ++ | ++ | +++ |
| *Cbp20*$^{KK109448}$/ *CG8888*$^{GD3777}$ | + | ++ | ++ | ++ | ++ | ++ | ++ | +++ | ++++ |
| *Cbp20*$^{KK109448}$/ *dlg1*$^{GD4689}$ | + | ++ | ++ | ++ | ++ | +++ | ++ | + | ++++ |
| *Cbp20*$^{KK109448}$/ *Fsn*$^{GD11383}$ | + | ++ | ++ | ++ | +++ | +++ | ++ | ++++ | ++++ |
| *Cbp20*$^{KK109448}$/ *Pak*$^{KK101874}$ | | ++ | ++ | + | ++ | ++ | + | + | +++ |
| *Cbp20*$^{KK109448}$/ *PIG-Z*$^{KK107404}$ | | | + | ++ | ++ | +++ | ++ | +++ | ++++ |
| *Overexp Diap1* | | | | | | | | | |
| *Cbp20*$^{KK109448}$/ *Overexp Diap1* | | | + | | ++ | + | | | |
| *dlg1*$^{GD4689}$/ *Overexp Diap1* | | | ++ | | | +++ | | | |
