## Supplementary material for "*NCBP2* modulates neurodevelopmental defects of the 3q29 deletion in *Drosophila* and *X. laevis* models": S9 Table

| Second-hit homolog | Cell cycle/ apoptosis | Microcephaly | ***Cbp20*** $^{KK109448}$ | ***CG8888*** $^{GD3777}$ | ***dlg1*** $^{GD4689}$ | ***Pak*** $^{KK101874}$ |
|---|---|---|---|---|---|---|
| *Arm (CTNNB1)* | X | | Enhancer (2/2) | Enhancer (2/2) | Enhancer (2/2) | Enhancer (2/2) |
| *Asp (ASPM)* | X | X | Enhancer (1/1) | Enhancer (1/1) | No interaction (0/1) | No interaction (0/1) |
| *Cadps (CADPS2)* | | | Enhancer (2/2) | Not validated (1/2) | Not validated (1/2) | Enhancer (2/2) |
| *Eph (EPHA6)* | | | Enhancer (3/3) | Enhancer (3/3) | Enhancer (3/3) | Not validated (1/3) |
| *kis (CHD8)* | X | | Not validated (1/2) | No interaction (0/2) | No interaction (0/2) | No interaction (0/2) |
| *Klp61F (KIF11)* | X | X | Enhancer (2/2) | Enhancer (1/1) | Enhancer (1/1) | Enhancer (1/1) |
| *MCPH1 (MCPH1)* | X | X | Enhancer (2/3) | Enhancer (3/3) | No interaction (0/3) | Not validated (1/3) |
| *nAChRα6 nAChRα7 (CHRNA7)* | | | Enhancer (3/5) | Enhancer (3/5) | Enhancer (3/5) | Enhancer (2/5) |
| *Nrx-1 (NRXN1)* | | | Enhancer (3/3) | Enhancer (3/3) | Enhancer (3/3) | No interaction (0/3) |
| *para (SCN1A)* | | | Enhancer (3/3) | Enhancer (2/3) | No interaction (0/3) | Not validated (1/3) |
| *Prosap (SHANK3)* | | | Not validated (1/2) | No interaction (0/2) | Not validated (1/2) | No interaction (0/2) |
| *Pten (PTEN)* | X | | Enhancer (2/2) | Enhancer (2/2) | Enhancer (2/2) | Not validated (1/2) |
| *Rk (LGR5)* | X | | Enhancer (4/5) | Enhancer (3/5) | No interaction (0/5) | No interaction (0/5) |
| *Sas-4 (CENPJ)* | X | X | Enhancer (2/2) | Enhancer (2/2) | Not validated (1/2) | No interaction (0/2) |
| *Ube3a (UBE3A)* | | | Enhancer (2/2) | Not validated (1/2) | No interaction (0/2) | Not validated (1/2) |
| **Lines tested (153)** | | | 39 | 38 | 38 | 38 |
| **All interactions (46/60)** | | | 15/15 | 13/15 | 9/15 | 9/15 |
| **Validated interactions (34/60)** | | | 13/15 | 11/15 | 6/15 | 4/15 |
