## Supplementary material for "*NCBP2* modulates neurodevelopmental defects of the 3q29 deletion in *Drosophila* and *X. laevis* models": S10 Table

| **3q29 deletion mouse models** | *B6J.Del16* $^{+/Bdh1\text{-}Tfrc}$ *(Baba et al.)* | *B6N.Del16* $^{+/Bdh1\text{-}Tfrc}$ *(Rutkowski et al.)* | *B6N.Dlg1* $^{+/-}$ *(Rutkowski et al.)* | *Pak2* $^{+/-}$ *(Wang et al.)* |
|---|---|---|---|---|
| Weight | Decreased | Decreased | No phenotype | Not tested |
| Brain size | Decreased | Decreased | Not tested | No phenotype |
| Locomotor activity | No phenotype | No phenotype | No phenotype | No phenotype |
| Amphetamine-induced locomotor activity | Not tested | Increased | Increased | Not tested |
| Anxiety (elevated plus maze or open field) | Not tested | No phenotype | No phenotype | No phenotype |
| Spatial learning and memory (water maze) | Not tested | Decreased | No phenotype | No phenotype |
| Acoustic startle response | Increased | Increased | No phenotype | No phenotype |
| Prepulse inhibition/sensorimotor gating | Decreased | No phenotype | No phenotype | No phenotype |
| Startle response w/risperidone | Rescued | Not tested | Not tested | Not tested |
| Marble burying | Not tested | No phenotype | No phenotype | Increased |
| Self-grooming | Increased | Not tested | Not tested | Increased |
| Social interaction (free or 3-chamber) | Decreased | Decreased | No phenotype | Decreased |
| Fear conditioning (context) | Decreased | No phenotype | No phenotype | Not tested |
| Auditory excitatory neuron activity | Increased | Not tested | Not tested | Not tested |
| Parvalbumin neuronal count | Decreased | Not tested | Not tested | Not tested |
| Dendritic spine density | Not tested | Not tested | Not tested | Decreased |
| Long-term potentiation | Not tested | Not tested | Not tested | Decreased |
| Synaptic density | Not tested | Not tested | Not tested | Decreased |
| Neuronal migration | Not tested | Not tested | Not tested | Decreased |
| Transcriptome | Immediate early signaling genes | Not tested | Not tested | Post-synaptic density, cytoskeleton, channel activity |
