## Supplementary material for "*NCBP2* modulates neurodevelopmental defects of the 3q29 deletion in *Drosophila* and *X. laevis* models": S11 Table

| **Candidate gene set** | **Overlap with apoptosis (%)** | **Simulated overlap with apoptosis** | | | **Percentile of observed overlap** | **Empirical p-value** |
|---|---|---|---|---|---|---|
| | | Min. | Mean | Max. | | |
| Autism (n=756) | 106 (14.0%) | 40 | 71 | 104 | 100% | $p<1.00\times10^{-5}$ |
| Intellectual disability (n=1,854) | 265 (14.3%) | 121 | 170 | 223 | 100% | $p<1.00\times10^{-5}$ |
| Schizophrenia (n=2,546) | 268 (10.5%) | 180 | 237 | 302 | 98.6% | p=0.014 |
