## Supplementary material for "*NCBP2* modulates neurodevelopmental defects of the 3q29 deletion in *Drosophila* and *X. laevis* models": S12 Table

| ***X. laevis* homolog** | **Morpholino** |
|---|---|
| *ncbp2* | for L, 5’- CGGTTTCCCTAGAATAGAAACAGGT-3’ |
| *fbxo45* | for L and S, 5’-TATCTGTGGTGGGAAGAAAAGGTCA-3’ |
| *dlg1* | for L, 5’-CAAATGAGGCAGCAACTTACTTTCT-3’ |
| *pak2* | for L and S, 5’-AGAGATAAATCCTACCTTTTTCTGT-3’ |
| standard control | 5′-cctcttacctcagttacaatttata-3′ |
