## Supplementary material for "*NCBP2* modulates neurodevelopmental defects of the 3q29 deletion in *Drosophila* and *X. laevis* models": S13 Table

| *X. laevis* homolog | Primers |
|---|---|
| *ncbp2* | forward for L allele 5’- ATCTGAGTCAGTATCGGGACC-3’<br>reverse for L allele 5’- CCCTTCCTTAAATCCTGCATCC-3’ |
| *fbxo45* | forward for L and S allele 5’- CCGACATACTGTGCAACCTG-3’<br>reverse for L and S allele 5’-TGTCCAAGATCACCCGAATCC-3’ |
| *dlg1* | forward for L allele 5’-CTCTCCTATGAACCCGTCAC-3’<br>reverse for L allele 5’-CCGGCCTCTATGAATTTGTG-3’ |
| *pak2* | forward for L and S allele 5’-AGGATAAACCACCAGCTCCTC-3’<br>reverse for L and S allele 5’-GGGAGCCCATCTTTATCTGGTG-3’ |
| *ODC1* control | forward 5’- GCCATTGTGAAGACTCTCTCCATTC-3’<br>reverse 5’- TTCGGGTGATTCCTTGCCAC-3’ |
